# gene2drug: a Computational Tool for Pathway-based Rational Drug Repositioning

**DOI:** 10.1101/192005

**Authors:** Francesco Napolitano, Diego Carrella, Barbara Mandriani, Sandra Pisonero, Francesco Sirci, Diego Medina, Nicola Brunetti-Pierri, Diego di Bernardo

**Affiliations:** Telethon Institute of Genetics and Medicine (TIGEM), Pozzuoli (NA), 80078, Italy; Department of Translational Medicine, Federico II University, 80131 Naples, Italy; Department of Chemical, Materials and Industrial Production Engineering, University of Naples Federico II, 80125 Naples, Italy

## Abstract

**Motivation:** Drug repositioning has been proposed as an effective shortcut to drug discovery. The availability of large collections of transcriptional responses to drugs enables computational approaches to drug repositioning directly based on measured molecular effects.

**Results:** We introduce a novel computational methodology for rational drug repositioning, which exploits the transcriptional responses following treatment with small molecule. Specifically, given a therapeutic target gene, a prioritisation of potential effective drugs is obtained by assessing their impact on the transcription of genes in the pathway(s) including the target. We performed in silico validation and comparison with a state-of-art technique based on similar principles. We next performed experimental validation in two different real-case drug repositioning scenarios: (i) upregulation of the glutamate-pyruvate transaminase, which has been shown to induce reduction of oxalate levels in a mouse model of primary hyperoxaluria, and (ii) activation of the transcription factor TFEB, a master regulator of lysosomal biogenesis and autophagy, whose modulation may be beneficial in neurodegenerative disorders.

**Availability:** free at http://gene2drug.tigem.it

## 1 Introduction

The study of approved drugs for new therapeutic applications, i.e. drug repositioning, is a potential shortcut in the drug discovery process [1]. Computational analysis of transcriptional responses of cells to chemical and genetic perturbations or in disease has been successfully applied for preclinical investigations of new drug indications [8, 4, 22, 11, 23, 16].

Transcriptomic approaches in industrial and academic settings leverage large scale collections of gene expression profiles such as the Connectivity Map (CMap [13]), which includes a total of 7,056 genomewide expression profiles obtained upon treatment of 5 different cell lines with different concentrations of 1309 small molecules. The CMap project is currently being scaled up by 3 orders of magnitudes by including 1.4M profiles, derived from treatment of 15 cell lines with 15,000 small molecules and 5000 genetic perturbagens, although only 1000 genes are measured in this case (LINCS [28, 18]).

Drugs inducing a transcriptional response opposite to the one induced by a disease may exert therapeutic effects independently of their molecular targets [4]. An advantage of transcriptomics approaches is that they can be applied in a completely data-driven fashion, without prior knowledge about disease or therapeutic mechanisms. Although fully data-driven approaches do not require any prior information, they cannot take advantage of it when available. From this perspective, a complementary well-established route to drug repositioning is provided by rational approaches. In rational drug repositioning, a specific therapeutic target gene is known in advance and drugs modulating its activity are investigated. Many drug-target prioritisation methods have been proposed in which chemo-structural considerations guide the selection of small molecules for their binding affinity with the target (target-based), or for their similarity to existing small molecules known to bind the target (ligand based) [15]. Nonetheless, 80% of newly discovered drugs tend to bind targets that are interactors of previously known therapeutic targets [5]. Available information about protein-protein interactions can thus be exploited to improve target-based drug discovery methods. Indeed, a number of computational drug repositioning approaches exploit the known protein interactors of the therapeutic target to predict the small molecules with highest probability of modulating the target [7, 30, 3].

In the context of rational approaches, transcriptional responses to drug treatment can also provide important information about drug mode of action, i.e. its molecular target [8, 19, 10]. However, drug-induced differential expression of the molecular target, if present, can be masked by the much larger differential expression of off-target genes [9]. Nevertheless, off-targets may be functionally related to the intended target, so that their differential expression level can be exploited as an indirect marker of the therapeutic target activity. For this reason, computational drug repositioning methods exploiting both drug-induced transcriptional responses and protein-protein interaction networks have been recently developed [12, 6, 9].

A recently published comparison of 13 different computational approaches to drug repositioning [9], including methods based on the expression of the target alone, on protein-protein interactions alone, or on a combinations of gene expression and proteinprotein interactions, found the best performance for a method of the latter type, namely “Local Radiality” (LR). LR takes into account the protein interactions among the significantly differentially expressed genes in the drug-induced transcriptional response and the therapeutic target in order to predict which drugs may modulate the target and thus are candidates for repositioning.

Here, we developed a novel approach to rational drug repositioning combining drug-induced transcriptional responses with annotated pathways as an alternative to protein interaction networks. Specifically, our method relies on the identification of drugs inducing significant transcriptional modulation of pathways that involve the target gene, as opposed to its protein interactors in a protein-protein interaction network. While this approach may prioritise drugs directly acting on the therapeutic target, any drug modulating the expression of the target-related pathways, even not directly, will be selected as a potential candidate for repositioning.

We implemented the method as on line tool named “Gene2Drug”, which takes advantage of publicly available pathway annotations from different sources. We computationally assessed the performance obtained by Gene2Drug using 10 different pathway databases and compared its performance both to the LR method and a naive method based on the target gene expression alone.

To investigate the efficiency of the method on real case scenarios, we tested Gene2Drug experimentally in two different settings: (i) to find drugs able to induce the expression of the Glutamic-Pyruvate Transaminase (GPT, aka Alanine Aminotransferase) whose over-expression was reported to reduce oxalate levels in mouse models of Primary Hyperoxaluria Type I, an inborn error of liver metabolism [21]; (ii) to find drugs activating the transcription factor TFEB, a master regulator of lysosomal biogenesis and autophagy, whose modulation maybe beneficial in the treatment of neurodegenerative disorders [24].

## 2 Methods

### 2.1 Approach

Gene2Drug uses gene expression data obtained from the Connectivity Map (CMap) [13], including genome-wide transcriptional response to treatments with 1,309 different small molecules. The CMap is currently the largest single collection of drug-induced gene expression profiles in which the expression of most genes is measured (LINCS data include the expression of just 1000 genes, while the expression of the other ≈11,000 genes is computationally inferred [28]). Gene2Drug relies on a pathway-based version of the CMap that we previously derived [19], in which all the pathways in a database are ranked according to how much the expression of genes annotated to each pathway changes after drug treatment, as shown in Fig. 1. Ten different pathway databases are supported (Tab. 1).

**Figure 1:**
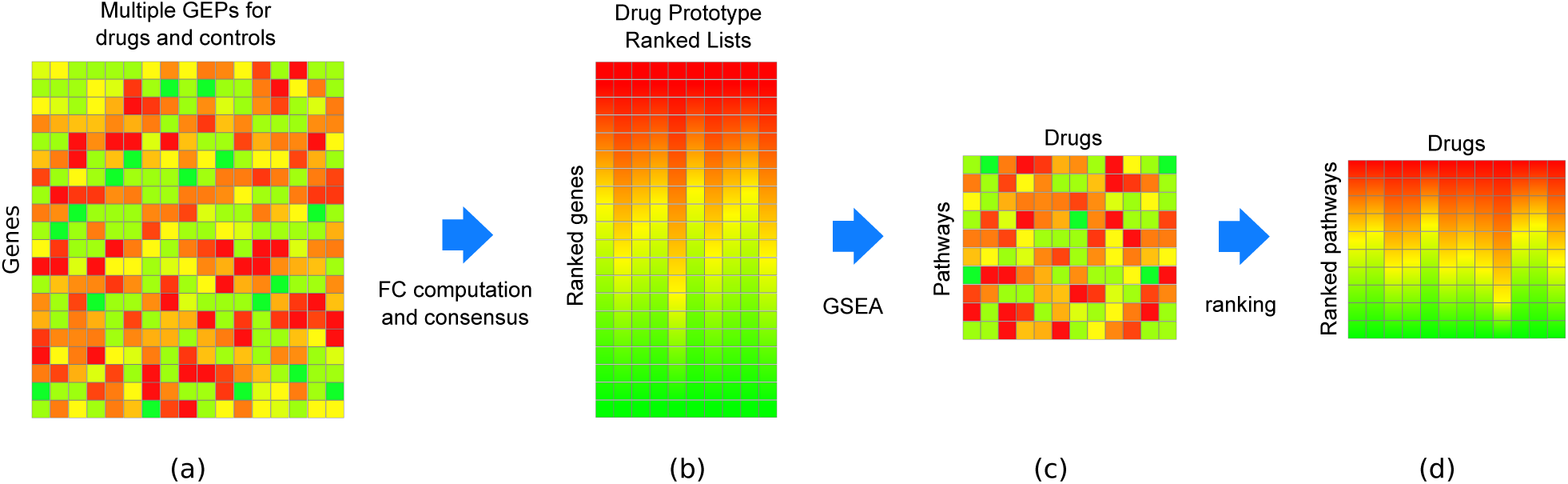
Bioinformatics pipeline to compute the pathway-based version of the Connectivity Map. (a) Raw genome wide expression profiles were collected from the Connectivity Map and preprocessed. (b) Control-treatment fold change values were computed and converted to ranks. Profiles referring to the same small molecule in different experimental conditions were merged together. (c) Enrichment Scores and p-values are computed for each Drug-pathway pair. (d) ESs are converted to column-wise ranks according to their p-values (most significantly upregulated on top, most significantly downregulated at the bottom).

**Table 1:**
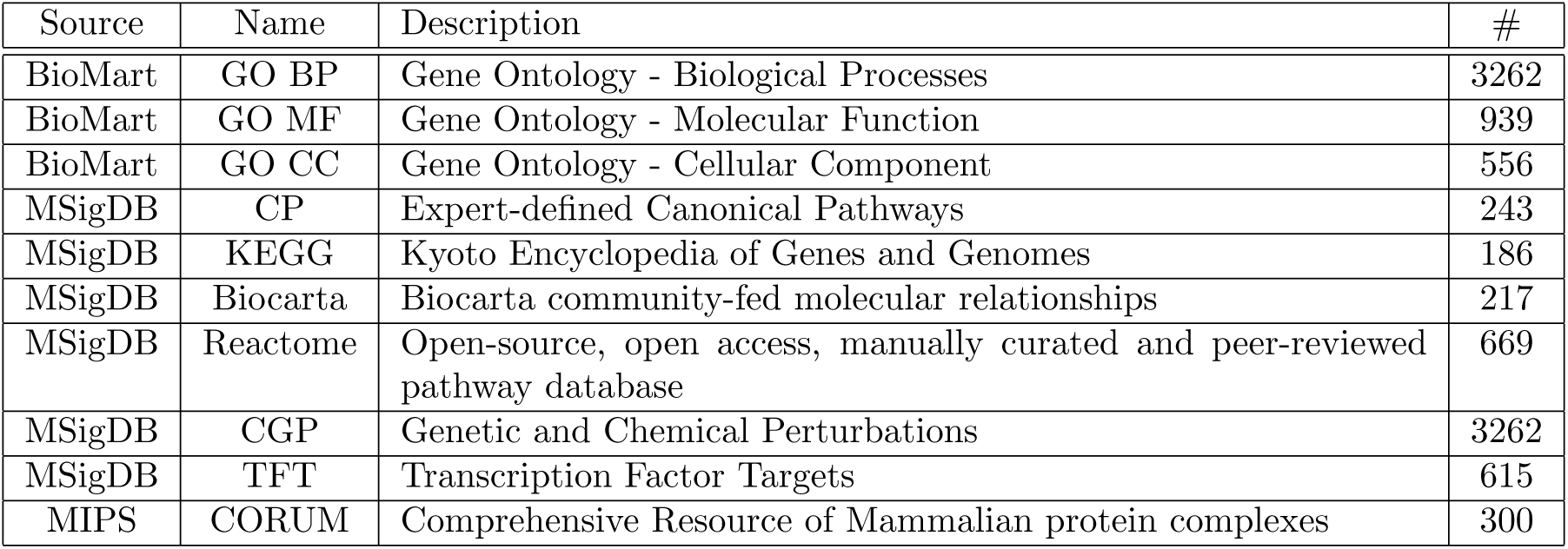
Pathway databases currently supported by Gene2Drug. Pathways were obtained from 10 publicly available collections.

Given a subset of pathways including the therapeutic target gene, Gene2Drug computes for each drug an Enrichment Score (ES) and its p-value according to how much they tend to be up- or downregulated by that drug, as shown in Fig. 2. This is done by applying Gene Set Enrichment Analysis (GSEA [25]) but for a set of pathways rather than a set of genes. Gene2Drug then outputs the list of 1,309 drugs ranked according to the computed p-values.

**Figure 2:**
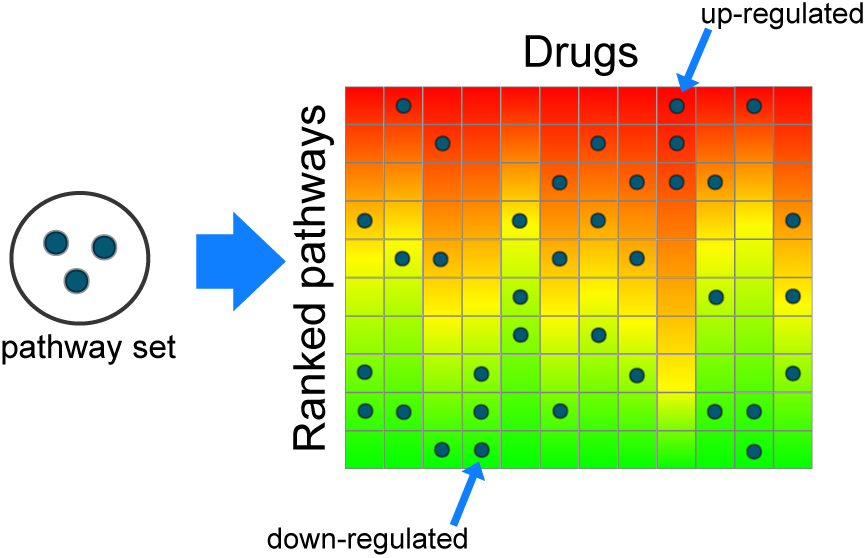
Schematics of the Gene2Drug approach. Given a set of pathways containing the therapeutic target, the Enrichment Score is computed for each drug to identify those able to significantly upregulate (or downregulate) the pathways in the set. In this example, the drug in the 4*th* column is predicted to inhibit the target, whereas the drug in the 9*th* column is predicted to activate the target.

To support gene-drug prioritisation directly, Gene2Drug takes a single gene as input and it generates automatically the subset of pathways including the input gene. While a user may want to manually select a specific subset of pathways of interest and possibly obtain better results in this way, the performance of the tool was assessed using this automatic selection.

### 2.2 Data preparation

The 6,100 differential gene expression profiles from the CMap were first reduced to 1,309 ranked lists of genes (one per drug), as shown in Fig. 1, by merging together those obtained with the same small molecule [8]. Each ranked list of genes was subsequently converted to a ranked list of pathways by means of Gene Set Enrichment Analysis (GSEA) [25]. GSEA uses a generalization of the Kolmogorov-Smirnov statistic as Enrichment Score (ES) to assess how much the genes in a pathway are distributed towards the top or bottom of the ranked list of genes. The actual implementation of GSEA used by Gene2Drug is available on line [19].

Given a database of pathways, we thus obtained a matrix of signed p-values (a negative p-value is the p-value of a pathway with a negative ES), with pathways along the rows and drugs along the columns, as shown in Fig. 1 [19]. Each element (*i, j*) of the ma trix thus contains a p-value assessing how significant the modulation of pathway *i* following treatment with drug *j* is. Finally, we ranked each of the columns according to the signed p-values, so that significantly up(down-) regulated pathways appear at the top (bottom) of each column (Fig. 1). We applied this procedure to 10 different pathway databases, listed in Tab. 1, thus obtaining 10 pathway-drug matrices.

### 2.3 Identification of drugs modulating a target gene of interest

Given a subset of pathways containing the target gene, Gene2Drug assesses how much these pathways tend to appear at the top or bottom of each ranked list of pathways (one for each drug), as exemplified in Fig. 2. To this end, GSEA is used to compute an Enrichment Score and a p-value for each drug. The final output is a list of drugs ranked by the corresponding p-values.

### 2.4 Experimental validation methods

Besides computational validation, Gene2Drug was tested in two different experimental settings. The corresponding experimental procedures follow.

#### 2.4.1 GPT: Luciferase assays

Both the human Huh-7 hepatic cells and the mouse Hepa1-6 hepatoma cells were cultured at 37 in Dulbecco’s modified Eagle’s medium (DMEM) supplemented with 10% fetal bovine serum (FBS) and 1% penicillin/streptomycin. Cells were plated in 24well plates and transfected with a reporter construct carrying the human GPT promoter driving the expression of the luciferase reporter gene (Switchgear Genomics) using Lipofectamine 2000 Transfection reagent according to the manufacturer’s instructions (Life Technologies). 24 hours post-transfection cells were treated with different concentrations of small molecule drugs: Fulvestrant (Sigma-Aldrich) (20*μM*, 50*μM*, 100*μM*, 125*μM*, 250*μM*, 500*μM*), Tomatidine (Extrasynthese) (5*μM*, 10*μM*, 20*μM*), and Nifuroxazide (Sigma-Aldrich) (5*μM*, 10*μM*, 20*μM*). Prior to the treatment, the concentrations of each small molecule were tested in cells and were not found to result in cell mortality by microscopic observation. DMSO was used as drug-vehicle. After 24 hours treatment, cells were washed with PBS, lysed and assayed for Renilla luciferase activity using the Dual-GLO Luciferase Assay System (Promega) by Glomax 96 microplate luminometer. All assays were performed in at least triplicate and the data are presented as means *±* standard deviation.

#### 2.4.2 TFEB: nuclear translocation assay

Following a previously described protocol [17], HeLa cells stably expressing TFEB-GFP construct were seeded in a 384-well plate, incubated for 24h and treated with the different compounds at 0, 0.1, 1, 10, 20 and 30 *μM* for additional 24h. After that cells were fixed with 4% paraformaldehyde or ice-cold methanol and permeabilized/blocked with 0.05% (w/v) saponin, 0.5% (w/v) BSA and 50 *μM* NH4Cl in PBS (blocking buffer). Images were acquired using the high content Opera system (Perkin Elmer) and analyzed using Harmony software and a dedicated script. Non-linear regression curves were determined by using Prism software.

## 3 Results

### 3.1 Gene2Drug

The aim of Gene2Drug is to identify one or more drugs able to modulate a therapeutic target of interest and thus proritise these drugs for repositioning.

As shown in Fig. 1, Gene2Drug makes use of publicly available data: the CMap, a collection of transcriptional responses to drug treatments, and a collection of annotated pathway databases, as listed in Tab. 1. Gene2Drug uses these data to build a ranked list of pathways for each drug in CMap representing the cellular response to drug treatment (Fig.1 and Methods). Pathways at the top (bottom) of the list include those genes which tend to be transcriptionally upregulated (downregulated) following drug treatment. This resource constitutes a higher level description of the well known ranked lists of genes and it is available for download from the Gene2Drug website. We derived 10 different versions of the pathway-wise CMap, one for each of 10 different pathway databases (Tab. 1) including signaling pathways, cellular components, biological processes, transcription factor targets, co-expressed and co-localized genes.

As shown in Fig. 2, and described in details in the Methods, Gene2Drug exploits the well-established GSEA statistics to rank the 1,309 drugs according to their ability to modulate the therapeutic target. To this end, Gene2Drug first identifies the set of pathways in the database that includes the therapeutic target. It then quantifies the drug-induced transcriptional modulation of these pathways by applying the GSEA method, where the gene-set is replaced by the pathway-set and the ranked list of genes by the ranked list of pathways. In this way, Gene2Drug assigns to each drug an Enrichment Score and a p-value. Finally, drugs are ranked according to their signed pvalue (a negative p-value is the p-value of a pathway with a negative Enrichment Score). Drugs at the top of the list are those predicted to most activate the therapeutic target, whereas the drugs at the bottom of the list, are the ones most inhibiting the therapeutic target.

Gene2Drug is implemented as a user-friendly web site publicly available at http://gene2drug.tigem.it. The web site supports both the manual input of a pathway set of interest and the automatic generation of the set starting from a target gene.

### 3.2 Validation

We first validated the method *in silico* in order to have a general assessment of its performances as compared to existing state-of-the-art tools. We then performed two experimental validations on different two molecular targets: the liver-specific enzyme GPT (aka ALT) and the transcription factor TFEB.

#### 3.2.1 In silico validation

To assess Gene2Drug performance in comparison to a state-of-the-art method, we implemented the LR method, which was reported to be the best performing one across 13 different approaches [9]. LR makes use of a protein-protein interaction (PPI) network as obtained from the STRING database [26]. Given the set of significantly differentially expressed genes (DEGs) following a drug treatment, the shortest paths across the PPI network from each DEG to the therapeutic target gene of interest is computed. The average length of such paths is used to score the drug-target pair (the shorter the better). Note that there is a weak correlation between LR and Gene2Drug as genes in the same pathway tend to be closer in the PPI network (Suppl. Fig. <u>S1</u>).

We also implemented a naive single-gene based method as a baseline to compare with. This method simply ranks drugs according to the differential expression of the therapeutic target of interest in the CMap dataset. We refer to this method as Single Gene Expression (SGE).

To build a gold standard, we followed the approach described by Isik et al [9] based on the STITCH protein-chemical database [27]. Of the 1,309 compounds in CMap, 607 small molecules are present in the STITCH database, corresponding to total of 133,146 drug-target pairs. This number includes 4 different evidence types (“experimental”, “prediction”, “database”, and “text-mining”). Each pair has a score for at least one of the four evidence types. A “combined score” is also provided, which is computed over the *non-missing* scores, but it is heavily biased by the “text-mining” evidence (61% of the pairs have a “text-mining” evidence, against 14%, 2%, and 33% respectively for the “experimental”, “prediction” and “database” evidences, see Suppl. Fig. <u>S2</u>). For this reason we used all the marginal scores separately. In order to retain the most reliable predictions for the gold standard, we selected only the drug-target pairs in the top quartile of the score. To limit the computational burden, when more than 5000 drug-target pairs are present in the top quartile, only the first 5000 were used (this condition happened only for the “database” and “text-mining” evidences).

Note that assessing Gene2Drug and other computational methods against such a gold standard is likely to provide significantly pessimistic outcomes, as they necessarily include a large number of false negatives. For example, it was noted that while the estimated number of possible drug targets lies between 6000 and 8000 [20, 29], only around 1300 known drugs are in DrugBank [14]. Nonetheless, the gold standard represents a fair common ground to assess the relative performances of different methods.

Given one of the 4 gold standards, we used Gene2Drug with each of the 10 pathway databases and compared the resulting performances with those obtained using the naive SGE and the state-of-theart LR methods. For the sake of clarity, we will refer to all of them as a set of 12 different methods.

Given one of the 12 methods, for each known target, we obtained a full weighed ranking of the 1,309 CMap small molecules (where the weights are given by the p-values for Gene2Drug, the gene ranks for the SGE, and the Local Radiality scores for the LR method). We then generated a single ranked list of drug-target pairs by merging together the lists obtained for the different targets according to their weights. Finally, we assessed the performance of each method by analysing how much the top ranked drugtarget pairs were enriched for true positives according to the 4 STITCH gold standards.

Figure 3 reports a summary of such analysis for the STITCH “experimental” gold standard which incluces only drug-target pairs supported by experimental evidence. In Fig. 3, the *precision* value (or Positive Predicted Values, PPV) is normalised against the expected PPV for a random ordering of drug-target pairs and it is plotted as a function of drug-target pairs ordered according to one of the 12 methods. For this gold standard, the Gene2Drug approach using the CC and TFT databases performed significantly better than the others, scoring up to 5 and 6 fold better then random respectively.

**Figure 3:**
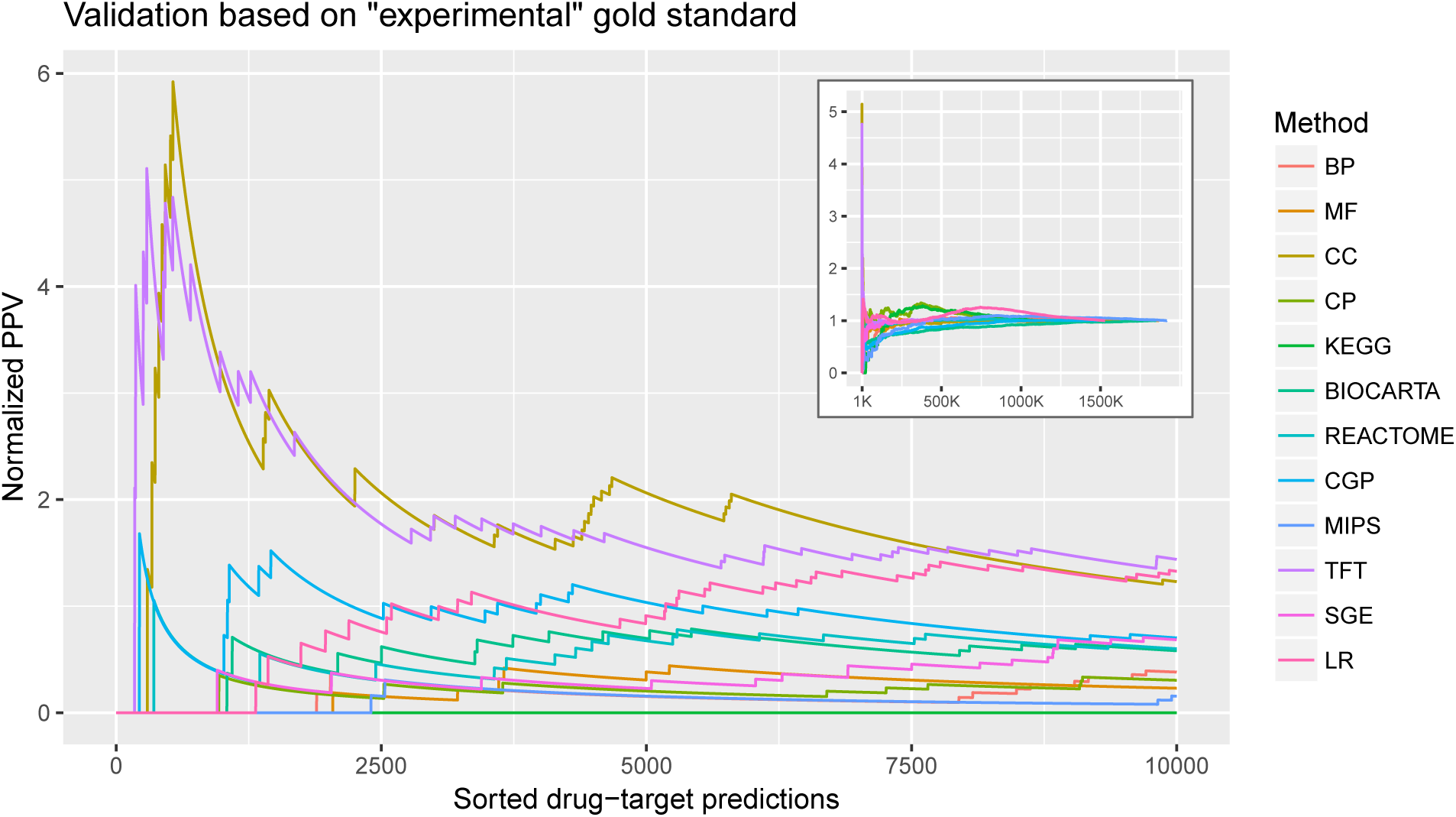
Validation of Gene2Drug against the STITCH experimental gold-standard. Only drugtarget pairs present in STITCH and supported by experimental evidence were included in this gold-standard (refer to Tab. 2 for additional gold-standards). For each one of the 12 methods, we obtained a list of drugtarget pairs ranked according to their weight as computed by the method. The Positive Predictive Value *TP/*(*TP* + *FP*) was computed for each rank, according to the STITCH experimental gold-standard, and divided by the “random” PPV obtained by ranking drug-target pairs randomly (Normalised PPV). Each line represents the PPV of one of the 12 methods as indicated in the legend for the top 10,000 drug-target pairs. **Inset:** PPV versus drug-targets but this time showing the PPV for all of the possible drug-target pairs, which vary according to the chosen database (BP: 1,853,544; MF: 1,864,016; CC: 1,233,078; CP: 1,117,886; KEGG: 1,127,049; BIOCARTA: 1,848,308; REACTOME: 1,424,192; CGP: 1,479,170; MIPS:1,916,376; TFT: 442,442; SGE: 335,104; LR: 1,524,985).

**Table 2:** Rankings of the methods across the 4 STITCH gold standards and corresponding medians. The Gene Ontology Cellular Component database (CC) showed the most consistent performance across all the gold standards except *Text-mining*. The Transcription Targets database (TFT) was the best performing method for the *Experimental* and *Prediction* gold standards. Local Radiality (LR) was the best for *Text-mining*. MIPS ranked top for *Database*, however performed poorly for *Experimental*, which is likely the most reliable gold standard. The Single-gene method (SGE) performed consistently worse than the others.

| Method | Experimental | Database | Prediction | Text-mining | Median |
| --- | --- | --- | --- | --- | --- |
| CC | 2 | 2 | 2 | 8 | 2.0 |
| TFT | 1 | 8 | 1 | 5 | 3.0 |
| LR | 3 | 4 | 4 | 1 | 3.5 |
| MIPS | 11 | 1 | 7 | 6 | 6.5 |
| MF | 7 | 7 | 10 | 7 | 7.0 |
| BIOCARTA | 6 | 10 | 8 | 3 | 7.0 |
| REACTOME | 5 | 3 | 9 | 12 | 7.0 |
| CP | 10 | 12 | 6 | 4 | 8.0 |
| CGP | 4 | 6 | 11 | 10 | 8.0 |
| BP | 8 | 9 | 3 | 9 | 8.5 |
| KEGG | 12 | 5 | 12 | 2 | 8.5 |
| SGE | 9 | 11 | 5 | 11 | 10.0 |

To further summarise the results, we computed also the median PPV of each method and for each gold standard considering only the top ranked 1% drug-target pairs. Tab. 2 reports the ranking of the methods according to this score for each gold standard. The Gene2Drug method with databases TFT and CC generally outperformed the LR method, which appeared particularly powerful when matched against the “text-mining” gold standards. The SGE method is almost always outperformed by the others, confirming again the hypothesis that the expression of the target gene alone is not a good predictor of drug mode of action.

#### 3.2.2. Experimental validation: upregulation of GPT expression

The glutamate-pyruvate transaminase (GPT) plays a key role in the intermediary metabolism of glucose and amino acids. GPT overexpression reduces oxalate in mouse models of Primary Hyperoxaluria Type I [21], a rare genetic disorder caused by loss-of-function mutations of the Alanine-Glyoxylate Aminotransferase gene (AGXT). We applied Gene2Drug to find drugs effective at increasing the expression of GPT to reduce the hyperoxaluria.

The GPT gene was annotated within 7 of our pathway databases. However, Reactome, a metabolism-centric database, included the pathways most relevant to the known GPT function (*metabolism of amino acids and derivatives* end *amino acid synthesis and interconversion transamination*). Note that this is an example of user-directed choice driven by prior knowledge on the therapeutic target function that is not readily supported by methods other than Gene2Drug.

We thus run Gene2Drug with the Reactome database using GPT as input. Table 3 shows the first 3 compounds (out of 1,309 compounds in CMap) ranked by Gene2Drug as those ones most upregulating the pathways involving GPT (a list of the top 30 is reported in Suppl. Tab. <u>S1</u>). We experimentally tested the efficacy of these 3 compounds to upregulate GPT in two different cell lines: Huh-7 (human hepatocytes) and Hepa1-6 (mouse hepatoma cells). Cells were transfected with the GPT promoter driving the expression of the luciferase reporter gene (Methods). Table 3 summarizes the results for the Huh-7 cells, additional details and results for the Hepa1-6 cells are reported in the supplemenatary materials. Fulvestrant resulted in significant upregulation of luciferase only at concentrations above 125*μM* (left panel of Fig. 4). Tomatidine resulted in dose-dendent increase of luciferase expression at low concentrations (right panel of Fig. 4). Nifuroxazide did not show any effect at concentration less than 20*μM* and was toxic in cells at higher concetrations (data not shown). Similar results for all the three compounds were confirmed also in the Hepa1-6 cell line (Suppl. Fig. <u>S3-S8</u>).

**Table 3:** Summary of luciferase assays for the three top ranked drugs predicted to upregulate GPT by Gene2Drug. A plasmid encoding for the luciferase reporter downstream of the GPT promoter was transiently transfected in Huh-7 cells (Methods). Each of the three compounds were tested at concentrations of 5*μ*M, 10*μ*M and 20*μ*M for 24 h. Fulvestrant did not show any effect at these concentrations, but it induced luciferase activity at higher concentrations (125*μM*, 250*μ* and 500*μM*) (Supplementary Information). Tomatidine significantly increased luciferase expression compared to vehicle control treatment and therefore higher concentrations were not tested (N.T.). Nifuroxazide failed to increase luciferase expression and higher concentrations were not tested because they resulted in cell toxicity.

| Rank | Compound | ES | p-value | Concentration<br>5-20μM | Concentration<br>125-500μM |
| --- | --- | --- | --- | --- | --- |
| **1** | **fulvestrant** | **0.98** | **0.002** | No | **Yes** |
| **2** | **tomatidine** | **0.97** | **0.002** | **Yes** | N.T. |
| **3** | nifuroxazide | 0.97 | 0.002 | No | toxic |

**Figure 4:**
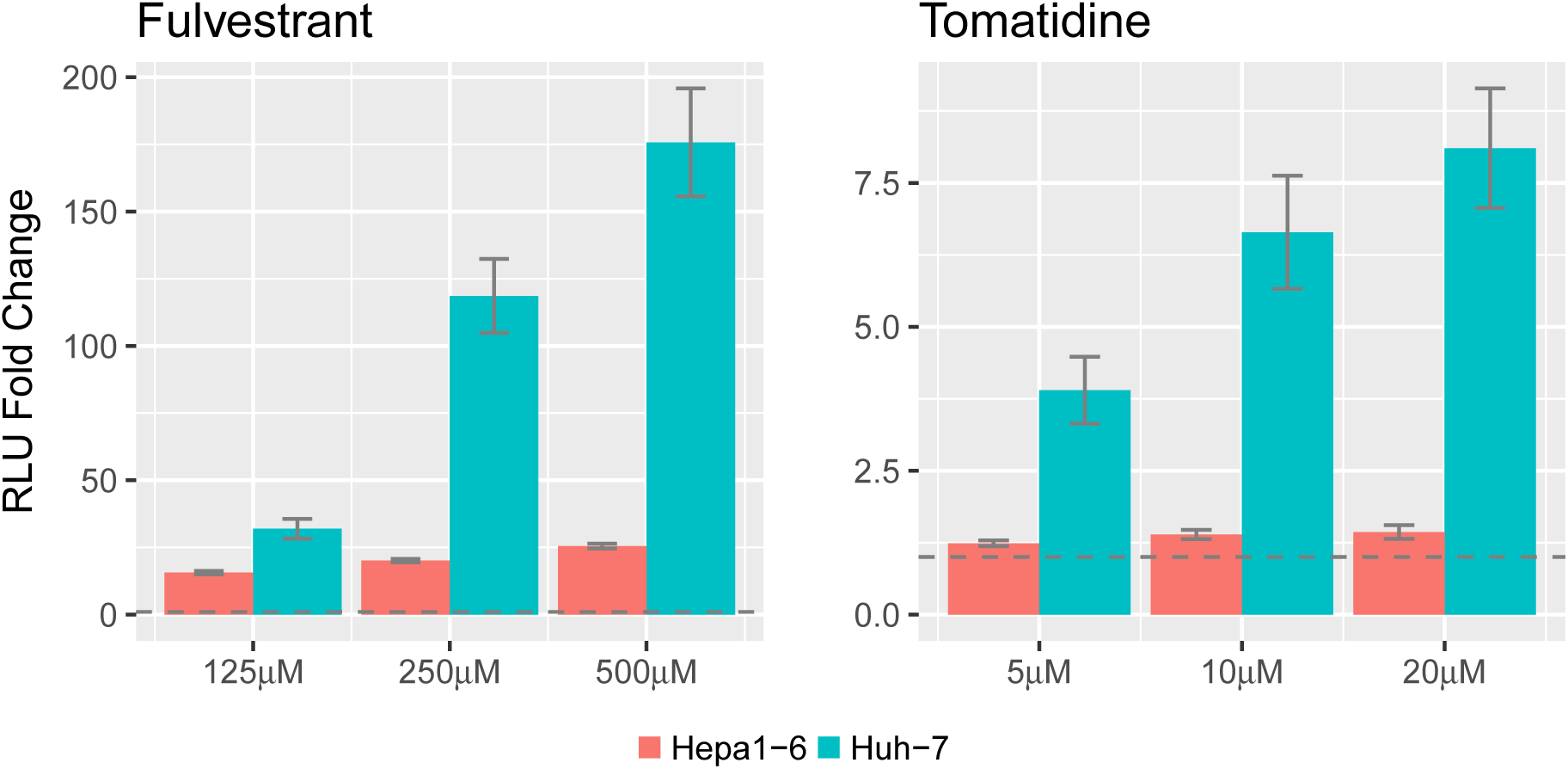
Experimental validation of predicted GPT modulators. Dose-dependent increase in relative luminescence units (RLU) in Hepa1-6 and Huh-7 cells transfected with a plasmid expressing the luciferase gene under the control of the GPT promoter and incubated with various concentrations of fulvestrant (left) and tomatidine (right). The dashed line idicates RLU fold change = 1 (no effect). The two compounds were ranked 1st and 2nd respectively among the small molecules predicted to upregulate pathways including GPT by Gene2Drug.

Because of the significant effect induced by fulvestrant and tomatidine, we wondered whether using the naive single gene expression (SGE) approach, i.e. simply ranking compounds according to the differential expression of GPT, would yield similar results. Surprisingly, SGE would rank fulvestrant 695th out of 1,309 among drugs overexpressing GPT, while GPT would be 6,126th out of 12012 among genes overexpressed by fulvestrant (data not shown). Similarly, tomatidine would rank 247th for GPT, while GPT would rank 2,391th for tomatidine (Suppl. Tab. <u>S2</u>).

#### 3.2.3 Experimental validation: induction of TFEB nuclear translocation

We then asked whether Gene2Drug can help identifying compounds able to modulate the activity of a transcription factor (TF) a particularly difficult task because TFs are usually considered undruggable targets [2]. To this end, we chose TFEB, a master regulator of lysosomal biogenesis and autophagy whose modulation has potential for the treatment of neurodegenerative disorders [24]. Also in this case, we chose to run Gene2Drug using a specific database based on the functional relevance of the pathways in which TFEB is annotated. We chose the *GeneOntology - Biological Processes* (GO-BP) database which included terms such as *lysosome organization* and *positive regulation of autophagy*.

Table 4 shows the 10 compounds (out of 1,309 compounds in CMap) ranked by Gene2Drug as those most upregulating the GO-BP terms containing TFEB.

**Table 4:** Drugs tested for TFEB nuclear translocation. 9 of the top 10 ranked by Gene2Drug were available to us. 4 of them induced TFEB translocation at concentrations lower than 30 *μ*M.

| Rank | Compound | ES | p-value | EC50 (μM) |
| --- | --- | --- | --- | --- |
| 1 | pimozide | 0.79 | 0.001 | > 30 |
| **2** | **deptropine** | **0.78** | **0.001** | **6.636** |
| **3** | **maprotiline** | **0.77** | **0.002** | **11.579** |
| 4 | nifurtimox | 0.75 | 0.002 | > 30 |
| 5 | benzethonium | 0.75 | 0.003 | > 30 |
| 6 | alprenolol | 0.75 | 0.003 | > 30 |
| 7 | 0297417-0002B | 0.74 | 0.003 | N.A. |
| **8** | **miconazole** | **0.74** | **0.003** | **7.223** |
| **9** | **loperamide** | **0.73** | **0.003** | **6.585** |
| 10 | etoposide | 0.73 | 0.003 | > 30 |

Among the top 10 drugs ranked by Gene2Drug, 9 were available to us. For each of the 9 drugs, we performed a High Content Screening assay for the TFEB nuclear translocation (TFEB-NT [17]) at 3h following drug administration at concentrations between 0.1*μ*M and 30*μ*M (Methods). Out of these 9 drugs, 4 were able to induce TFEB nuclear translocation, 3 of which at concentrations below 10 *μ*M, as reported in Tab. 4. A representative experimental result for deptropine (one of the 4 positives drugs) is shown in Fig. 5.

**Figure 5:**
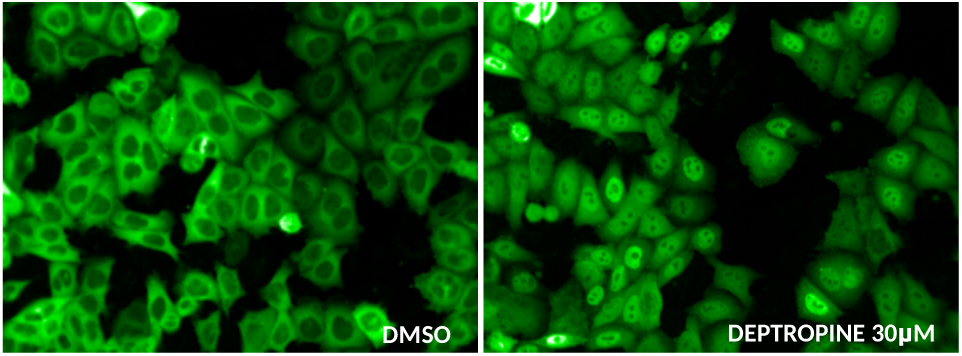
Deptropine induces TFEB nuclear translocation. Hela cells stably expressing TGEBGFP were seeded, incubated for 24h and treated with deptropine 30 *μ*M or DMSO 0.2% (control) for 6h.

## 4 Discussion and Conclusions

We introduced a computational approach for rational drug repositioning integrating transcriptional responses to small molecules with prior knowledge in the form of annotated pathway databases.

Gene2Drug implements a complementary approach to other state-of-the-art computational methods which exploit prior knowledge in the form of gene and protein interaction networks [9].

Gene2Drug is designed to be a semi-automated pipeline where the user chooses the pathways that best describe the function of the target gene, whose pharmacological modulation is deemed to be therapeutic.

Gene2Drug ranks small molecules in the CMap database according to their ability to induce transcriptional changes both in the therapeutic target as well as in the other genes in the selected pathways.

Using the STITCH database as a gold standard for drug-chemical interactions, we demonstrated that Gene2Drug consistently outperforms the naive singlegene method, where drugs are ranked according to the differential expression of the therapeutic target gene only.

We experimentally validated Gene2Drug in two different settings: targeting a metabolic enzyme (GPT) and a Transcription Factor (TFEB). We showed that in both cases, Gene2Drug was effective at identifying small molecules with the desired effects, at least in cell lines, whose known direct targets where either unknown (e.g. tomatidine) or completely unrelated to the desired effect (e.g. deptropine).

Gene2Drug can be easily extended to larger collections of gene-expression profiles, such as the new LINCS database. However the L1000 platform, which LINCS is based on, actually measures the expression of ˜1000 genes, with all the others being computationally inferred. While methods based on the similarity of transcriptional responses may not be significantly impacted by this limitation, the effectiveness of pathway enrichment analysis on L1000 data remains to be investigated.

In conclusion, Gene2Drug’s approach to rational drug repositioning combines transcriptomics with prior knowledge in the form of pathway databases, and it is complementary to those methods based on protein interaction networks.

